# Geldanamycin inhibits Jumonji histone lysine demethylase KDM4 and targets chimeric transcription factor PAX3-FOXO1

**DOI:** 10.1101/2019.12.11.872838

**Authors:** Ahmed Abu-Zaid, Shivendra Singh, Wenwei Lin, Jonathan Low, Alireza Abdolvahabi, Bailey Cooke, John Bowling, Sivaraja Vaithiyalingam, Jie Fang, Duane Currier, Mi-Kyung Yun, Alaa AlTahan, Dinesh M. Fernando, Julie Maier, Heather Tillman, Purva Bulsara, Zhaohua Lu, Sourav Das, Zhenmei Li, Brandon Young, Richard Lee, Zoran Rankovic, Stephen White, Andrew M. Davidoff, Taosheng Chen, Jun Yang

## Abstract

Histone lysine demethylases (KDMs) are emerging as therapeutic targets in cancer. Development of potent KDM inhibitors may provide additional options for epigenomics-oriented therapies. Using a Time-Resolved Fluorescence Resonance Energy Transfer (TR-FRET) functional demethylation assay, in combination with a high-content immunofluorescence imaging phenotypic screen, Matrix-Assisted Laser Desorption/Ionization-Fourier Transform Ion Cyclotron Resonance mass spectrometry (MALDI-FTICR MS) and Amplified Luminescent Proximity Homogeneous Assay (ALPHA), we identified geldanamycin, an inhibitor of heat shock protein 90 (Hsp90), as a novel inhibitor of JmjC-domain containing demethylases such as KDM4B. We further found that geldanamycin can destabilize the PAX3-FOXO1 fusion oncoprotein, an Hsp90 client, which is a driver of clinically unfavorable alveolar rhabdomyosarcoma (aRMS). We then hypothesized that dual inhibition of PAX3-FOXO1 and epigenetic modifiers of aRMS would have synergistic antitumor activity. We repurposed the geldanamycin analog 17-DMAG to target aRMS and found that 17-DMAG significantly delays tumor growth, extends survival in xenograft mouse models, and inhibits expression of PAX3-FOXO1 targets and multiple oncogenic pathways including MYC, E2F and NOTCH. In addition, the combination of 17-DMAG with conventional chemotherapy or the bromodomain inhibitor JQ1 significantly enhances therapeutic efficacy. In summary, we have identified geldanamycin and 17-DMAG as dual KDM/Hsp90 inhibitors and 17-DMAG is efficacious against PAX3-FOXO1-driven rhabdomyosarcoma.

## Introduction

The heat shock pathway plays a significant role in promoting protein folding. It is activated in many tumors and provides transformed cells with a survival advantage by maintaining protein homeostasis. The heat shock protein 90 (Hsp90) is harnessed by cancer cells to facilitate the function of oncoproteins as a molecular chaperone^1,2^; hence, one important feature of cancer cells is their “addiction” to Hsp90^3^, making this a potential therapeutic vulnerability. Since the identification of the first prototype of Hsp90 inhibitor^4^, geldanamycin, numerous Hsp90 inhibitors have been developed and at least seventeen entered clinical trials^5^, including the geldanamycin analogs 17-AAG (17-N-allylamino-17-demethoxygeldanamycin) and 17-DMAG (17-dimethylaminoethylamino-17-demethoxygeldanamycin). However, Hsp90 inhibitors have not demonstrated significant clinical efficacy^6^. The inhibitory potency and affinity of geldanamycin and its analogs for the isolated Hsp90 protein was in the low micromolar range^7-9^, which is in contrast to their low nanomolar cellular antiproliferative activity^8-11^, suggesting that the antiproliferative activity may be due to a mechanism other than via Hsp90 inhibition alone. A better understanding of the mechanisms of action of these Hsp90 inhibitors that underlie treatment response is key to improving their therapeutic activity and clinical outcomes.

Aberrant histone lysine methylation is commonly seen in a variety of cancers^12^, due to genetic alteration or dysregulated expression of histone lysine methyltransferases and histone lysine demethylases (KDMs)^13-18^. The KDM4 (KDM4A-D) subfamily are Jumonji-domain containing KDMs, which are responsible for removing methyl groups from tri- and dimethylated H3K9 and H3K36^19^. KDM4B is particularly important and is involved in a variety of pathophysiological functions^20^. We and others have shown that KDM4B is a direct target of estrogen receptor alpha (ERα) and hypoxia-inducible factor 1 (HIF1) in ERα^+^breast cancer^21-24^, and that it epigenetically regulates G2/M phase cell cycle gene expression^24^. In addition, KDM4B is a key molecule in androgen receptor (AR) signaling^25^, which epigenetically enhances AR transcriptional activity. Recently, we further showed that KDM4B is involved in neuroblastoma growth and tumor maintenance^26^. KDM4 members are also required for the transformation of pediatric leukemia driven by the fusion oncoprotein MLL–AF9^27,28^. A recent study has shown that KDM4B is involved in the regulation of unfolded protein response (UPR) in *PTEN*-deficient triple-negative breast cancers, and genetic depletion or small molecule inhibition of KDM4B activates the UPR pathway, resulting in preferential apoptosis^29^. These data suggest that targeting KDM4B may enhance the efficacy of proteotoxic drugs such as Hsp90 inhibitors by further impairing oncoprotein folding.

Rhabdomyosarcoma (RMS) is a devastating myogenic cancer in children, adolescents and young adults^30,31^. This soft tissue sarcoma is mainly classified into two histological subtypes, alveolar RMS (aRMS) and embryonal RMS (eRMS). aRMS is more aggressive, with a higher rate of metastasis and a poorer prognosis^30,32,33^. aRMS is primarily driven by the pathognomonic fusion oncoprotein PAX3-FOXO1^34,35^ or its variant PAX7-FOXO1. Although current treatment modalities have steadily improved survival of RMS patients, the outcome for aRMS patients with metastatic disease remain dismal, underscoring the pressing need to develop novel therapies for this subset of patients.

We recently identified ciclopirox as a new KDM4B inhibitor^36^, which shows antitumor activity in neuroblastoma models. Here we further found that ciclopirox induces differentiation of aRMS cells and inhibits tumor growth in PAX3-FOXO1-positive aRMS xenografts, indicating that pharmacologically targeting KDM may have therapeutic benefit to aRMS, which promoted us to develop more potent KDM4B inhibitors. By using multiple orthogonal validation approaches, we identified the Hsp90 inhibitors, geldanamycin and its analog 17-DMAG^4,37^, as novel and potent KDM4 inhibitors. We also found that PAX3-FOXO1 is an Hsp90 client, which was destabilized by geldanamycin. We therefore hypothesized that repurposing these ansamycins may achieve better efficacy against PAX3-FOXO1-driven aRMS. Indeed, 17-DMAG significantly delayed tumor growth. RNA-seq analysis showed that 17-DMAG affected multiple oncogenic pathways. Immunohistochemical staining showed that angiogenesis was significantly reduced while cell death markers (Caspase 3 and TUNEL) were significantly increased by 17-DMAG. The combination of 17-DMAG with conventional chemotherapy or the bromodomain inhibitor JQ1 further enhanced the antitumor efficacy in mouse xenograft models. These data indicate that dually targeting KDM and Hsp90 in aRMS by 17-DMAG provides a new paradigm for repurposing this old drug in targeted cancer therapy.

## Results

### TR-FRET demethylation assay identifies geldanamycin as a potent KDM4 inhibitor

We developed a primary 384-well format screening, Time-Resolved Fluorescence Resonance Energy Transfer (TR-FRET) demethylation functional assay, to identify novel KDM4B inhibitors (**Fig. 1A**). The assay uses a Terbium (Tb)-labeled anti-H3K9me2 antibody as a fluorescence donor, and an AF488 tagged streptavidin as a fluorescence acceptor that detects the biotinylated histone H3K9me2 peptide. When uninhibited, KDM4B converts the substrate H3K9me3 peptide to the product H3K9me2 peptide, which is recognized and bound by both the donor Tb-labelled antibody and acceptor AF488-labelled streptavidin. The resulting proximity of the Tb donor and the AF488 acceptor elicits a fluorescence emission at 520 nm when excited at 340 nm. When the KDM4B activity is inhibited, less biotin-H3K9me2 is generated and the 520 nm emission signal (and the 520 nm/490 nm ratio) is reduced. We optimized the conditions by (1) confirmation of the specificity of the Tb-labeled anti-H3K9me2 antibody to the Biotin-H3K9me2 peptide among the 4 relevant peptides (Biotin-H3K9me0, Biotin-H3K9me1, Biotin-H3K9me2 and Biotin-H3K9me3), (2) confirmation of the specificity of Tb-anti-H3K9me2 antibody to the product Biotin-H3K9me2 peptide over the substrate Biotin-H3K9me3 peptide over a wide concentration range (0.3 nM to 312.5 nM), (3) optimization of the KDM4B concentrations, (4) optimization of incubation time, (5) optimization of KDM4B activity in three selected buffers, (6) optimization of the salt types and their concentrations in buffers, and (7) evaluation of DMSO tolerance of the assay (**Supplementary Fig. 1A-1H**). Our TR-FRET assay displayed predictability and reproducibility of responses to known KDM4 inhibitors and showed a clear threshold between positive and negative responses (**Fig. 1B, 1C**). The high-throughput screening (HTS) statistical parameter Z’(Z-prime) had an average value of 0.73 (0.55-0.87) from our pilot screen (**Fig. 1D**), indicating that the assay is robust and reproducible. On the basis of the optimized parameters, we performed a pilot screening of 3262 FDA-approved drugs and bioactive molecules and identified the Hsp90 inhibitor geldanamycin as a novel and potent KDM4B inhibitor with an IC_50_ of 50.1 nM (**Fig. 1E, 1F**).

**Figure 1.**
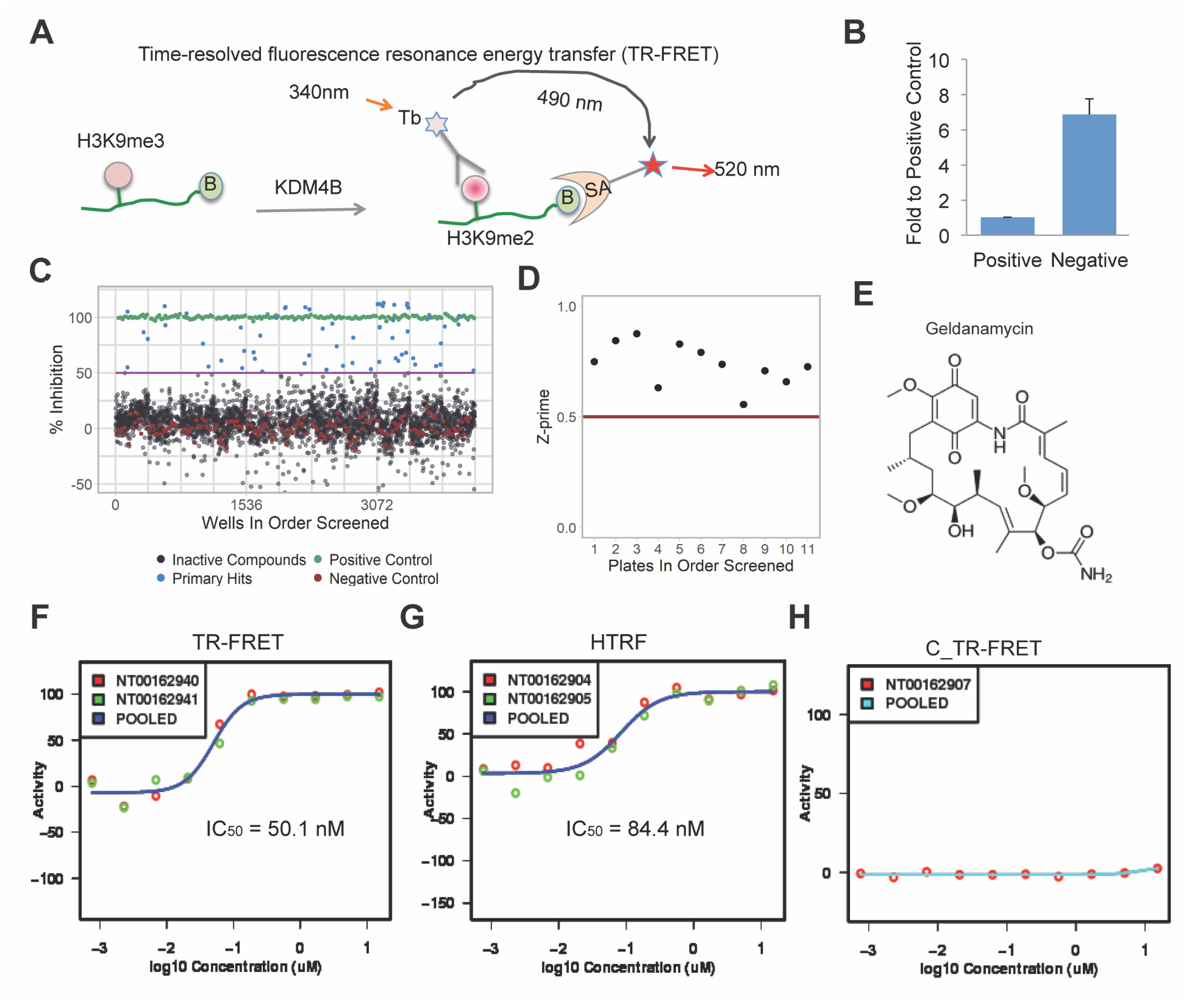
TR-FRET demethylation assay identifies geldanamycin as a potent KDM4 inhibitor. **(A)** The KDM4B TR-FRET demethylation functional assay: As a KDM4B substrate, the Biotin-linked H3K9me3 peptide is converted to the product Biotin-H3K9me2 peptide, which is bound by both the Terbium-labeled anti-H3K9me2 antibody and AF488-streptavidin and brings the donor fluorophore Tb and acceptor fluorophore AF488 into close proximity. A high TR-FRET signal in the form of light emission at 520 nm from the acceptor fluorophore AF488 is generated when the donor fluorophore Tb was excited at 340 nm. (B =biotin, SA = streptavidin, Tb = Terbium) **(B)** Fold ratio between positive and negative controls from the validation assay. The positive control is the group without KDM4B protein (mimic 100% inhibition) and the negative control is the group with 750 nM in-house KDM4B protein (mimic 0% inhibition). **(C)** Boxplot of activity values from the screen. **(D)** Z-prime factor from the screen. **(E)** The chemical structure of geldanamycin. **(F)** Dose-response curve of geldanamycin to KDM4B assayed by TR-FRET. **(G)** Dose-response curve of geldanamycin to KDM4B assayed by HTRF. **(H)** Background TR-FRET dose response curve of geldanamycin without KDM4B protein.

We also designed a homogeneous time-resolved fluorescence resonance energy transfer assay (HTRF) that is similar to TR-FRET, but the fluorophore AF488 is replaced with AF647, which switches the light detection wavelength from 520 nm to 665 nm to avoid interference by intrinsically fluorescent compounds and reduce false positives. The HTRF assay obtained a comparable IC_50_ value (84.4 nM) for geldanamycin (**Fig. 1G**). The inhibition of KDM4B by geldanamycin appears to be KDM4B-mediated because omitting KDM4B abrogated the drug response relationship (**Fig. 1H**). To test whether other non-ansamycin Hsp90 inhibitors were able to inhibit KDM4B activity, we included additional Hsp90 inhibitors (Ganetespib, KW2478, SNX-5422, AT13387, NVP-AUY922, STA-4783 and XL888); none of them showed an activity >15% of inhibition to KDM4B (**Supplementary Fig.2**). These data suggest geldanamycin, but not other non-ansamycin Hsp90 inhibitors, inhibits KDM4B.

### Orthogonal identification of geldanamycin as a KDM4 inhibitor by high-content immunofluorescence imaging screen and a MALDI-FTICR mass spectrometry-based approach

We also developed an orthogonal phenotypic assay in parallel with the TR-FRET assay as a secondary screen. We made a retroviral vector to transfer the wild type KDM4B gene and a catalytically dead mutant of KDM4B (H189A), in which histidine 189 was replaced by alanine, resulting in the loss of iron binding activity, leading to the loss of demethylation function. We then transduced U2OS cells with these retroviral vectors (**Fig. 2A**). In our experience, many cell lines cannot tolerate overexpression of KDM4B, due, perhaps, to a DNA damage response to global loss of H3K9me3, but U2OS cells appeared to tolerate the overexpression and exhibited the expected global reduction of H3K9me3 by the wild-type KDM4B (**Fig. 2A**). We then used these cell lines to screen 2684 compounds and monitored the H3K9me3 levels with immunofluorescence imaging in 384-well plates. The Z-prime score reached 0.5 (data not shown), indicating that this assay is acceptable for screening. We again found that geldanamycin inhibited KDM4B, with an IC_50_ of 21.2 nM in the cell-based assay (**Fig. 2B, 2C**). We further assessed the effect of geldanamycin and its analog 17-DMAG on KDM4B activity in U2OS cells via Western blot analysis (**Fig. 2D**). Both compounds blocked KDM4B catalytic activity on its H3K9me3 and H3K36me3 substrates in cells but not on H3K4me3, which is not a substrate of KDM4B (**Fig. 2D**). We performed a similar experiment in 293T cells expressing KDM4B or the catalytically dead mutant and included ciclopirox as a control. Geldanamycin, 17-DMAG, and ciclopirox greatly inhibited KDM4B activity in the cells expressing KDM4B, compared to those expressing the control or catalytically dead mutant (**Supplementary Fig. S3**). These data indicate that both geldanamycin and its analog 17-DMAG inhibit KDM4B activity in cells. To further validate the on-target of geldanamycin, we developed a Matrix-Assisted Laser Desorption/Ionization-Fourier Transform Ion Cyclotron Resonance (MALDI-FTICR) mass spectrometry-based approach, which is detailed in the Methods section. Again, we found that geldanamycin inhibited KDM4B activity with an IC_50_ of 1.24μM (**Fig. 2E, 2F**). Although this method was less sensitive than our TR-FRET and immunofluorescence assays, it validated that KDM4B activity was inhibited by geldanamycin in vitro.

**Figure 2.**
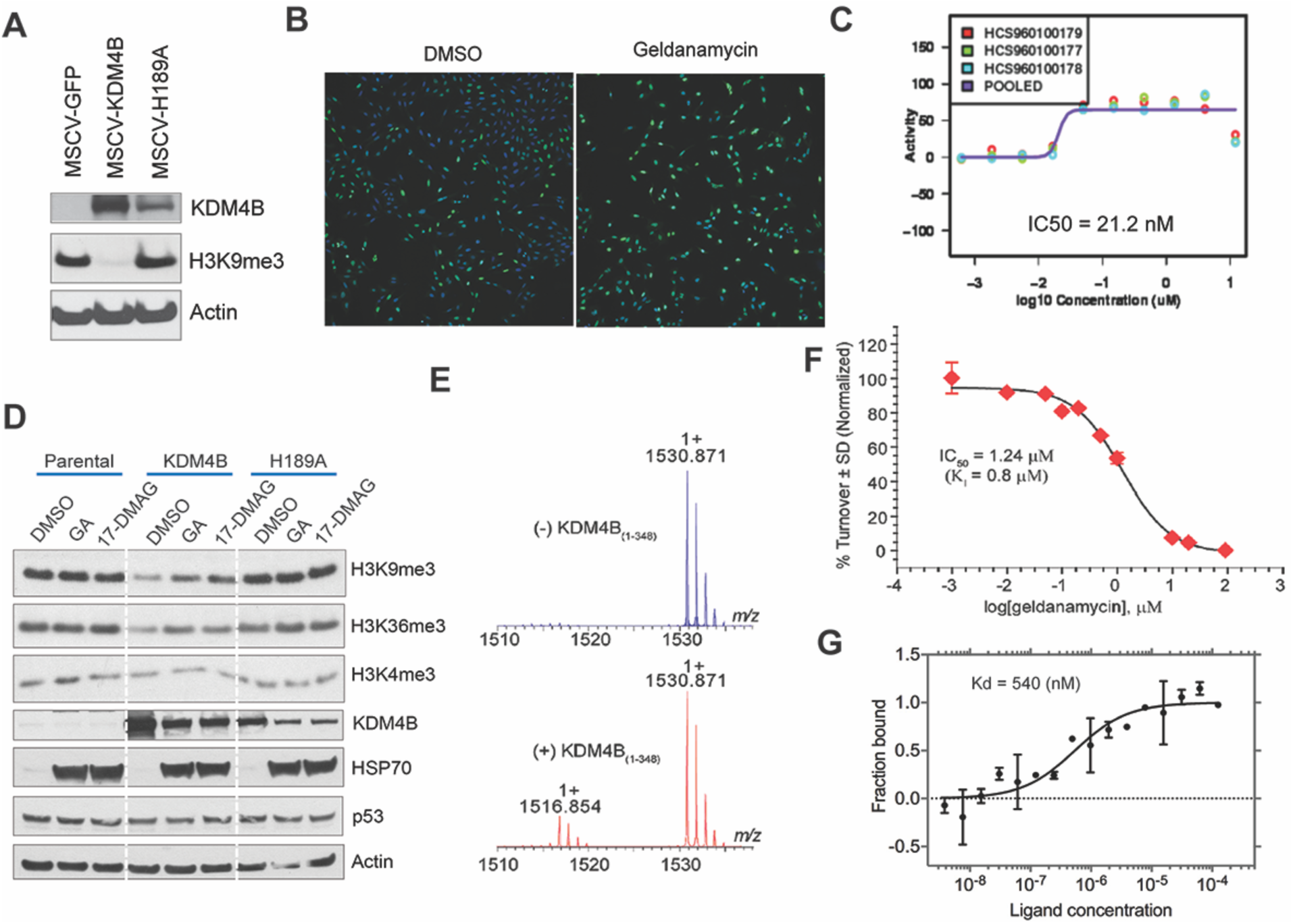
Orthogonal identification of geldanamycin as a KDM4 inhibitor by high-content immunofluorescence imaging screen and a MALDI-FTICR mass spectrometry-based approach. **(A)** Western blot for KDM4B expression in U2OS cells. **(B)** High-content immunofluorescence assay shows geldanamycin inhibits KDM4B activity. Green = H3K9me3, Blue = DAPI. **(C)** Dose-response curve shows IC50 of KDM4B inhibition by immunofluorescence assay. (**D**)Western blot shows 1uM of geldanamycin and its analog 17-DMAG block KDM4B activity in U2OS cells, as evidenced by changes in histone methyl marks assessed by specific antibodies. **(E)** MALDI-FTICR mass spectrometry shows that KDM4B converts H3K9me3 to H3K9me2. **(F)** Dose-response curve of geldanamycin shows the inhibition of KDM4B with IC50 of 1.24 uM. **(G)** Microscale Thermophoresis assay of direct binding of KDM4B and geldanamycin. Kd = dissociation constan

To determine the binding affinity of geldanamycin and the catalytic domain of KDM4B, we performed a Microscale Thermophoresis (MST) assay, which showed a Kd value of 540 nM (**Fig. 2G**). These data indicate a direct binding of geldanamycin and the catalytic domain of KDM4B.

### Profiling of geldanamycin activity against KDM using ALPHA screen

To determine the selectivity of geldanamycin and its analogs 17-DMAG and 17-AAG against KDMs, an ALPHA screen was used to assess the KDM inhibitory activity of these ansamycins at 5 μM. Geldanamycin showed greatest inhibition to KDM4 and KDM5 subfamilies (**Fig. 3A**), which have highest sequence homology in comparison to other KDMs with a phylogenetic tree showing that KDM4 and KDM5 members are the closest neighbors (**Fig. 3A**). These data suggest the inhibitory activity of geldanamycin is correlated with JmjC domain structures. Interestingly, 17-DMAG had a broader inhibitory activity to KDMs including KDM3-6 (**Fig. 3A**). In contrast, 17-AAG showed a high inhibition to KDM4A and KDM4C (**Fig. 3A**). We further determined the IC_50_ of the three ansamycin analogs to KDMs, and we chose 1-2 KDMs that were representatives of each subfamily. Geldanamycin has an IC_50_ below 1μM to KDM4B, KDM4C and KDM5A (**Fig. 3B**), while 17-DMAG had an IC_50_ below 1μM to KDM3A, KDM4B, KDM4C, KDM5A and KDM6B (**Fig. 3B)**. However, 17-AAG had an IC_50_ below 1μM to KDM4C and KDM5A (**Fig. 3B)**. All three compounds were much less effective at inhibiting KDM1A (IC_50_>10μM), a flavin adenine dinucleotide-dependent amine oxidase domain containing KDM, which is structurally distinct from JmjC KDMs. All three compounds showed a reasonable dose-response relationship (**Fig. 3C, 3D**). These data indicate that the inhibitory activity of ansamycins to KDMs depends on the JmjC domain and is impacted by the modifications at 17-position of the benzoquinone ring of the compounds (**Fig. 3E**).

**Figure 3.**
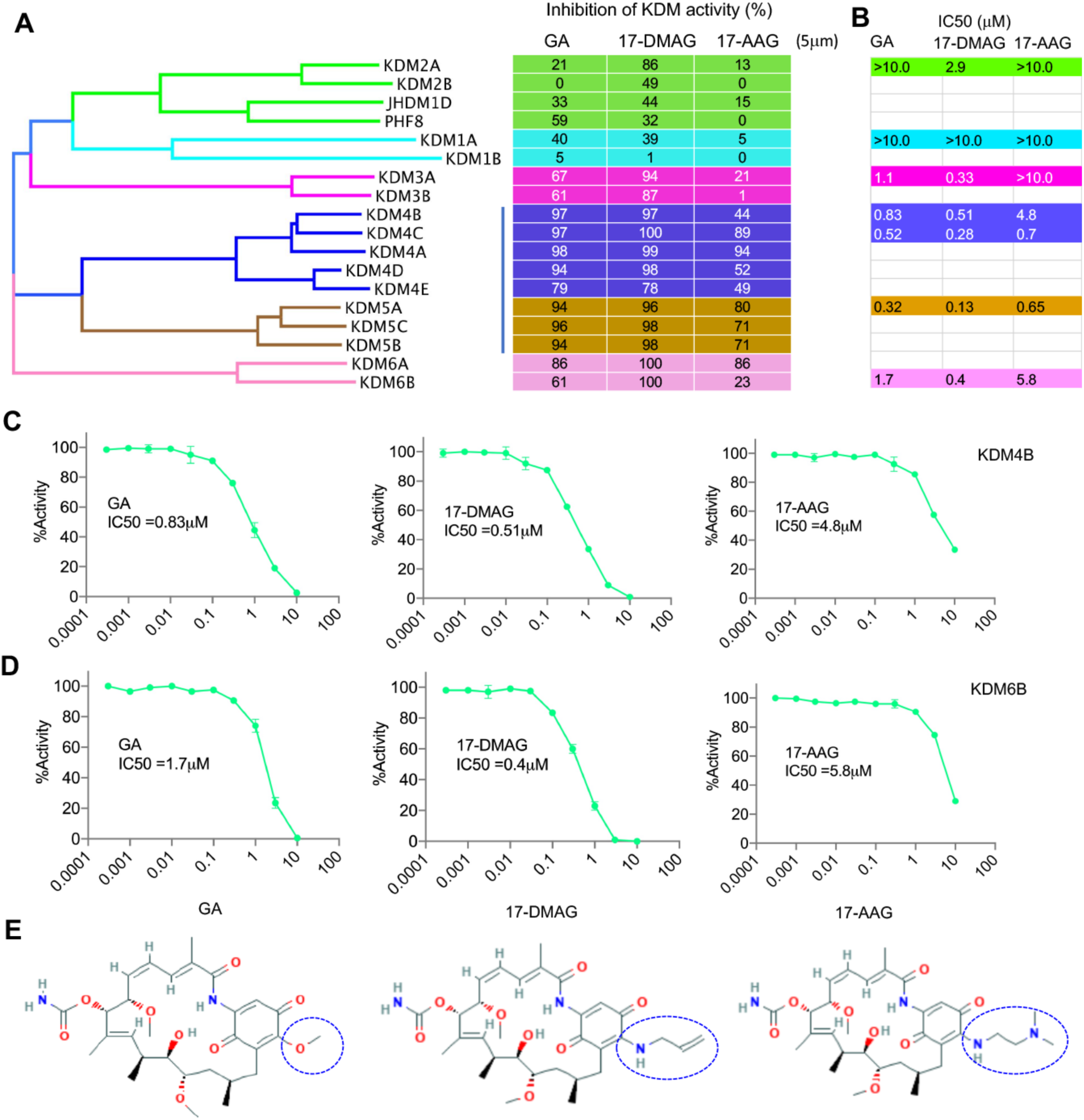
Profiling of geldanamycin activity against KDM using ALPHA screen. **(A)** Phylogenetic tree of KDMs (left) and KDM selectivity profiling of geldanamycin, 17-DMAG and 17-AAG at 5μM (right) assayed by ALPHA Screen. **(C)** IC50 of geldanamycin, 17-DMAG and 17-AAG for selected KDMs assayed by ALPHA Screen. (**D**) Dose-response curve of geldanamycin, 17-DMAG and 17-AAG against KDM4B and KDM6B. (**E**) 17-position of benzoquinone moieties of Geldanamycin, 17-DMAG and 17-AAG are circled (structures were obtained from Pubchem, https://pubchem.ncbi.nlm.nih.gov)^56^.

### Geldanamycin promotes PAX3-FOXO1 degradation in aRMS cells

As an oncogenic transcription factor, PAX3-FOXO1 is difficult to target directly. Previous studies have shown a chaperone dependency of fusion oncoproteins such as BCR-ABL^38^, EML4-ALK^39^ and EWS-FLI1^40^, as they are Hsp90 clients and are destabilized by Hsp90 inhibitors. Since chimeric oncoproteins do not exist in normal cells, we hypothesized that fusion proteins such as PAX3-FOXO1 may be more prone to degradation in the absence of cellular chaperone protein function. We examined whether PAX3-FOXO1 expression was downregulated by geldanamycin and 17-DMAG, known Hsp90 inhibitors. Immunoprecipitation confirmed that PAX3-FOXO1 and Hsp90 physically complexed (**Fig. 4B**). Indeed, PAX3-FOXO1 expression was reduced by geldanamycin and 17-DMAG (**Fig. 4A**). Inhibition of Hsp90 often leads to proteasomal degradation of its clients^6^. Consistent with this, the downregulation of PAX3-FOXO1 by geldanamycin or 17-DMAG was rescued by MG132 (**Fig. 4C**), a reversible proteasome inhibitor. These data demonstrate that the PAX3-FOXO1 protein needs Hsp90 for its stabilization, providing the rational to target PAX3-FOXO1 protein stability by inhibiting Hsp90 activity. Thus, geldanamycin not only inhibits the enzymatic activity of KDM but also downregulates key oncoprotein levels by targeting Hsp90, thus making it a unique dual inhibitor.

**Figure 4.**
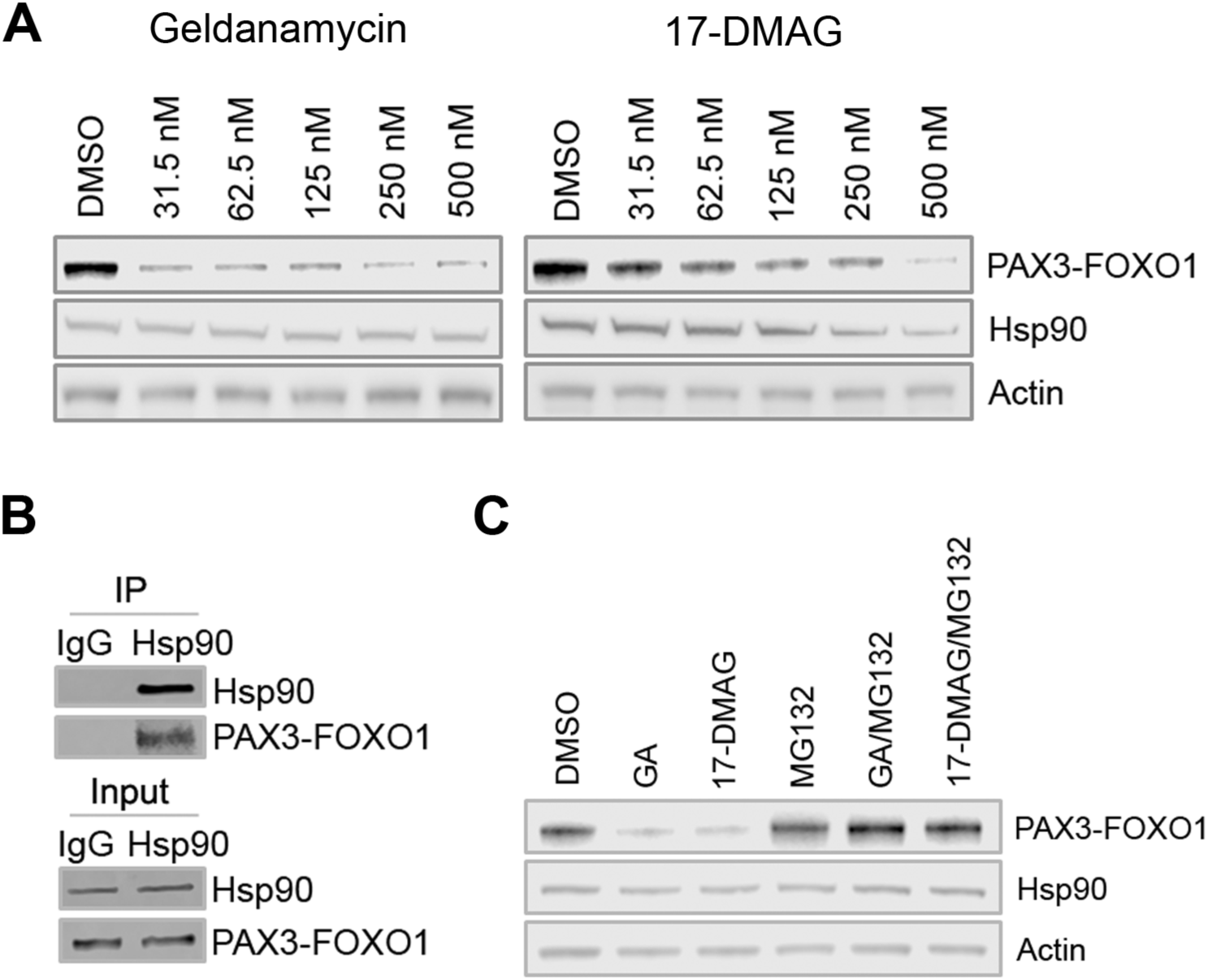
Geldanamycin promotes PAX3-FOXO1 degradation in aRMS cells. **(A)** After 24-hour treatment with geldanamycin or 17-DMAG, Rh30 cells were lysed for western blotting with the indicated antibodies. **(B)** Immunoprecipitation of Rh30 cell lysates with mouse IgG or monoclonal Hsp90 antibody, followed by western blotting with indicated antibodies. **(C)** Rh30 cells were treated with 200nM of geldanamycin or 17-DMAG with or without 5μM of MG132 for 24 hours. Western blot was performed with the indicated antibodies.

### 17-DMAG suppresses tumor growth, inhibits tumor angiogenesis, and disrupts multiple oncogenic pathways

We previously identified ciclopirox as a KDM4 inhibitor that has antitumor activity in neuroblastoma models^36^. We have further shown that ciclopirox induced RMS cell differentiation and inhibited tumor growth in a PAX3-FOXO1-positive aRMS xenograft model (**Supplementary Fig. 4**). We have also determined that KDM4 inhibition is a vulnerability of PAX3-FOXO1-driven RMS using a selective KDM4 inhibitor QC6352^41^ (manuscript in preparation), which epigenetically impacts expression of PAX3-FOXO1 targets. We therefore hypothesized that geldanamycin/17-DMAG might be effective for PAX3-FOXO1-positive aRMS based on: (1) their ability to inhibit KDM4 activity; and (2) their ability to target PAX3-FOXO1 for proteasomal degradation. As geldanamycin has unfavorable pharmacokinetics in vivo and is associated with liver toxicity, we chose 17-DMAG and 17-AAG for in vivo assessment. After PAX3-FOXO1-positive Rh30 xenografts implanted in CB17 *scid* mice reached about 200 mm^3^ in size, 17-DMAG or 17-AAG was given intraperitoneally at a dose of 25mg/kg or 50mg/kg, respectively, twice daily, every 4 days. 17-DMAG treatment significantly delayed tumor growth (**Fig. 5A**) and extended mouse survival (**Fig. 5B**). In contrast, 17-AAG had only a very modest effect on tumor growth and survival (**Fig. 5A, 5B**). RNA-seq analysis of the treated xenografts showed that PAX3-FOXO1 targets such as *FGFR4*, were significantly downregulated by 17-DMAG but not 17-AAG (**Fig. 5C**). FGFR4 is known to be important for RMS tumor growth^42^. Gene set enrichment analysis (GSEA) showed that 17-DMAG significantly inhibited MYC, E2F and NOTCH pathways (**Fig. 5D**), all of which are essential to cancer cell growth, proliferation and survival. Immunohistochemical staining for the apoptosis marker Caspase 3 and a cell death marker TUNEL showed that 17-DMAG induced significant cancer cell death (**Fig. 5E**). In addition, the areas of blood vessels indicated by angiogenesis marker CD31, which labeled endothelial cells, were greatly reduced (**Fig. 5F**), indicating that 17-DMAG inhibits tumor angiogenesis. Taken together, 17-DMAG has potent antitumor effect and targets multiple oncogenic pathways in aRMS.

**Figure 5.**
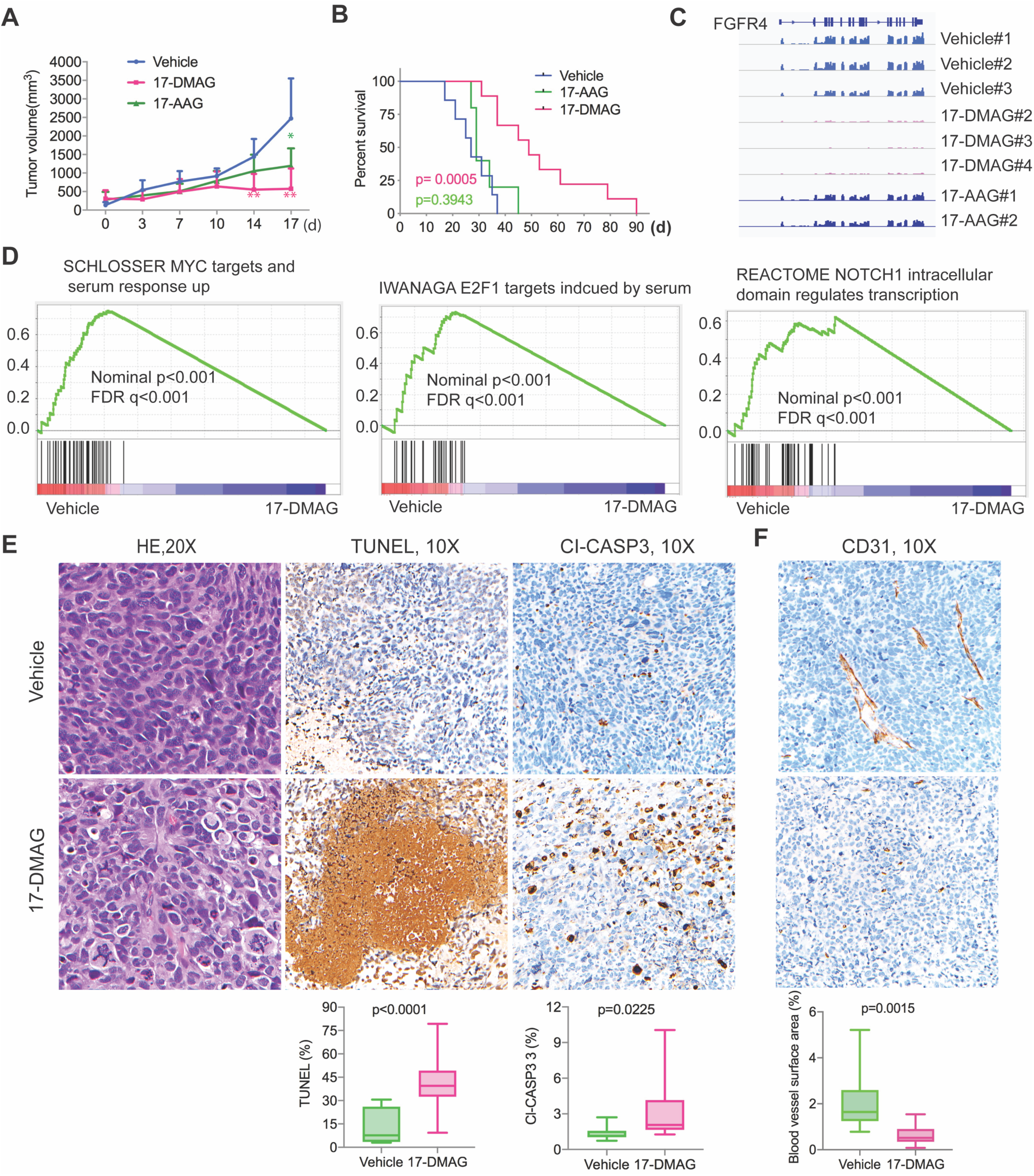
17-DMAG suppresses tumor growth, inhibits tumor angiogenesis, and disrupts multiple oncogenic pathways. (A) Tumor growth curve of Rh30 in CB17 *scid* mice treated with vehicle (n=7), 25mg/kg of 17-DMAG (n=11) and 50mg/kg of 17-AAG (n=7). Unpaired t test for comparison of tumor volumes of each group. **p<0.01, *p<0.05. **(B)** Kaplan-Meier analysis of mouse survival treated with vehicle, 17-DMAG and 17-AAG. Log-rank test for comparison of survival of each group. **(C)** RNA-seq read of *FGFR4* from 3 individual tumors of vehicle, 3 individual tumors of 17-DMAG and 2 individual tumors of 17-AAG. (**D**) GSEA analyses of pathways downregulated by 17-DMAG. **(E)** Hematoxylin and Eosin (H&E), cleaved caspase 3 (Cl-CASP3) and TUNEL immunohistochemistry staining of tumor tissue sections from vehicle and 17-DMAG treatment. 3 different areas per section from 4 tumors (n=12) in each group were compared with unpaired student t test. **(F)** CD31 immunohistochemistry staining of tumor tissue sections from vehicle and 17-DMAG treatment.

### Combination of 17-DMAG with conventional chemotherapy or bromodomain inhibitor JQ1 enhances therapeutic efficacy

A recent preclinical study showed that combination of the Hsp90 inhibitor Ganetespib with Vincristine (VCR) and Irinotecan (IRN) significantly enhanced the tumor response in RMS xenograft models^43^. Considering that Ganetespib alone showed no significant effect on inhibition of RMS xenograft^43^ while 17-DMAG significantly delayed tumor growth (**Fig. 5A**), we anticipated that the combination of 17-DMAG and VCR+IRN may obtain better efficacy. Indeed, combining 17-DMAG significantly improved the efficacy of VCR/IRN in two PAX3-FOXO1-positive RMS xenograft models implanted in NSG mice (**Fig. 6A-6F)**. In the Rh30 xenograft model, all mice had complete response (CR) to the combination therapy while nearly all control mice had progressive disease (PD) when treated with 17-DMAG alone or chemotherapy alone (**Fig. 6A-6C**). In the Rh41 xenograft model, combination therapy also showed better response (11% stable disease, 22% partial response and 67% complete response) than monotherapy (**Fig. 6D-6F**).

**Figure 6.**
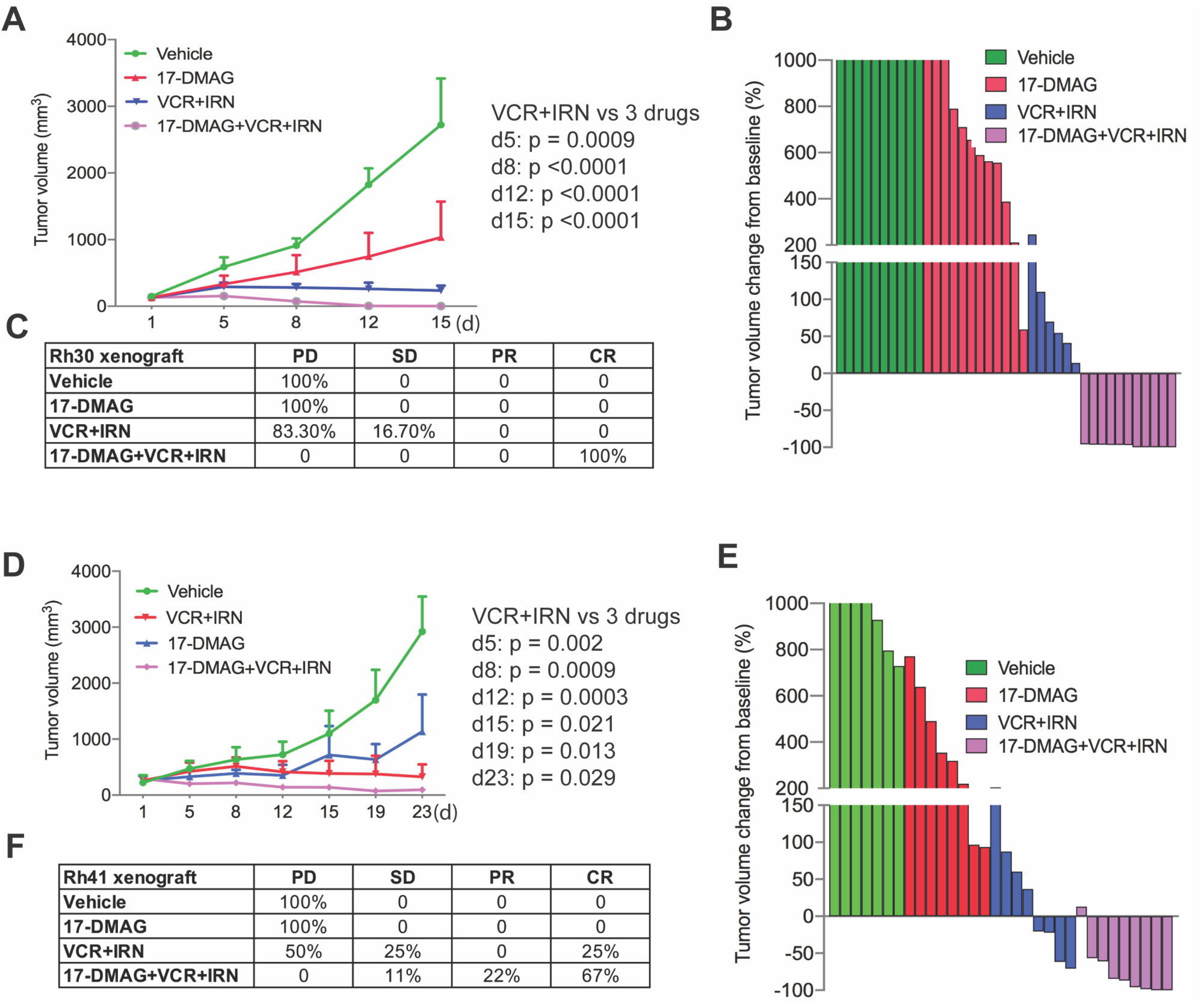
Combination of 17-DMAG with conventional chemotherapy enhances therapeutic efficacy. (**A-C**) Tumor growth curve of Rh30 in NSG mice treated with vehicle (n =10), 17-DMAG (n=12), VCR/IRN (n=6), 17-DMAG/VCR/IRN (n=11) for two weeks (**A**), Waterfall plot of response to treatment with 17-DMAG, VCR/IRN, and 17-DMAG/VCR/IRN (**B**), Summary of tumor response to treatment with 17-DMAG, VCR/IRN, and 17-DMAG/VCR/IRN (**C**). *p* value is determined by Wilcoxon Rank Sum test between treatment groups VCR+IRN and 17-DMAG+VCR+IRN. (**D-F**) Tumor growth curve of Rh41 in NSG mice treated with vehicle (n =7), 17-DMAG (n=8), VCR+IRN (n=8), 17-DMAG+VCR+IRN (n=9) for three weeks (**D**), Waterfall plot of response to the treatment with 17-DMAG, VCR+IRN, and 17-DMAG+VCR+IRN (**E**), Summary of tumor response to the treatment with 17-DMAG, VCR/IRN, and 17-DMAG/VCR/IRN (**F**). *p* value is determined by Wilcoxon Rank Sum test between treatment groups VCR+IRN and 17-DMAG+VCR+IRN.

Finally, based on a recent study that showed PAX3-FOXO1 was dependent on BRD4 and that the bromodomain inhibitor JQ1 suppressed tumor growth^44^, we hypothesized that the combination of 17-DMAG and JQ1 would synergistically enhance the therapeutic efficacy. In a PAX3-FOXO1-positive Rh41 xenograft model, the single agent modestly delayed tumor growth, while the combination significantly reduced tumor growth (**Supplementary Fig. 6A-6C**). Taken together, 17-DMAG showed potent anti-tumor activity when combined with conventional therapy or targeted therapy.

## Discussion

Geldanamycin, and its analog 17-AAG and 17-DMAG, are prototype Hsp90 inhibitors which bind N-terminal pocket of Hsp90^45^ and have potent anticancer activity. However, the mechanism of action of these ansamycins is not entirely clear. It has been reported that geldanamycin and its analogs have inhibitory activity and binding affinity to Hsp90 in the range of 0.3–10 μM^7-9^, which is in contrast to the low nanomolar antiproliferative activity of the compounds in multiple cell lines in culture^8-11^. Three major mechanisms were proposed to interpret the discrepancy of the 100-fold greater potency in cell culture. The first theory is ansamycins bind to and inhibit an Hsp90 multiprotein complex with much higher affinity than to Hsp90 alone^46^. The second explanation is that the physicochemical properties of the ansamycins result in its intracellular accumulation from cell culture media^9^, leading to highly potent antiproliferative activity. The third possible reason may be due to geldanamycin’s time-dependent, tight binding to Hsp90^47^. However, in this study, we identified geldanamycin and 17-DMAG as potent KDM inhibitors using multiple orthogonal validation approaches (TR-FRET, high-content immunofluorescence, MALDI-FTICR mass spectrometry, alpha screen and MST assay), convincingly demonstrating that these compounds are epigenetic modulators. Other chemotypes of Hsp90 inhibitors showed no direct KDM inhibition, indicating that geldanamycin has unique features being a dual inhibitor of KDM/Hsp90.

KDMs play an important role in carcinogenesis, metastasis and therapy resistance^48^. Pharmacologically targeting KDM effectively inhibits tumor growth in multiple preclinical models. A selective KDM1 inhibitor has entered clinical trial in Ewing sarcoma treatment. We recently identified an antifungal drug ciclopirox as a JmjC-domain containing KDM inhibitor, which targets KDM4B and suppresses neuroblastoma growth^36^. Here, we further showed this compound inhibits rhabdomyosarcoma growth, providing a proof-of-concept that KDM is a potential vulnerability of solid tumors driven by oncogenic transcription factors such as MYC and PAX3-FOXO1, whose activity may require epigenetic modifiers to facilitate their transformation activity. Considering that most oncogenic transcription factors are difficult to target directly as they usually do not bear druggable pockets, targeting such epigenetic modifiers may provide alternative options to inhibiting cancer drivers. The surprising discovery of geldanamycin and its analogs as KDM inhibitors suggests that cancers driven by oncogenic transcription factors may be more sensitive to geldanamycin and 17-DMAG than other chemotypes of Hsp90 inhibitor, considering that solid tumors are hypoxic and multiple *KDMs* are hypoxia-inducible genes^49^. While the chaperone dependency of oncoproteins is already known, dependency of PAX3-FOXO1 on Hsp90 for protein stability has not been previously reported. Our study shows that PAX3-FOXO1 physically interacts with Hsp90 and pharmacologic inhibition of Hsp90 induces proteasome-dependent degradation of PAX3-FOXO1, indicating that PAX3-FOXO1 is a new Hsp90 client. These data provide evidence that the first prototype Hsp90 inhibitor geldanamycin and its analog 17-DMAG are new KDM inhibitors, exerting dual inhibition of Hsp90 and histone demethylases, which provides a rational for the repurposing of 17-DMAG in cancer treatment. 17-DMAG showed potent antitumor activity and inhibited PAX3-FOXO1 targets such as FGFR4. Multiple oncogenic pathways including MYC, E2F1 and NOTCH were inhibited. Previous studies have shown that MYC and NOTCH pathway play an important role in rhabdomyosarcoma^50-52^. In addition, we found that 17-DMAG has anti-angiogenic effect, which is consistent with a previous finding^53^. Combination of 17-DMAG with conventional chemotherapy further enhanced efficacy. With the rationale that BRD4 also engages in PAX3-FOXO1-mediated function, we further extended our hypothesis that combination of 17-DMAG with BRD4 inhibitor would enhance therapeutic efficacy. Indeed, combination of 17-DMAG with the bromodomain inhibitor JQ-1 remarkably inhibited tumor growth compared with monotherapy. These data indicate that mechanism-based rational combination therapy may achieve a better antitumor efficacy. As chimeric transcription factors such as MLL-AF9 and MOZ-TIF2 play a critical role in cellular transformation, and need KDM4 for their function^26,28^, geldanamycin analogs might be suitable to target cancers driven by such oncofusion proteins.

Geldanamycin is a natural antibiotic isolated as the fermentation product of *Streptomyces hygroscopicus*, a bacterial strain widely distributed in nature, especially in the soil. Activity profiling of geldanamycin against KDMs showed that geldanamycin is more selective to inhibit human KDM4 and KDM5 members, two closest subfamilies in the KDM phylogenetic tree (**Fig. 3A**). Why would a bacterium produce an antibiotic that is able to dually inhibit KDM and Hsp90? One naïve speculation is that it may help bacteria more effectively defend surrounding competitors such as other bacterial strains or fungi. Although bacteria do not have homologs of KDM4 and KDM5, they do have JmjC domain containing proteins and Hsp90 homologs. However, fungi have homologs of KDM4 and KDM5 based on the phylogenetic analysis (**Supplementary Fig. 5**). 17-DMAG and 17-AAG, two derivatives of geldanamycin used in clinical application, have distinct selectivity against KDMs. While 17-DMAG showed a broader KDM inhibition to KDM3, KDM4, KDM5, and KDM6, 17-AAG seemed to be more selective to KDM4A and KDM4C. This might be the reason that 17-DMAG had a higher therapeutic efficacy than 17-AAG in suppressing tumor growth in aRMS xenograft models. Particularly, all three compounds showed very low activity to KDM1A (IC_50_>10μM), which is not a JmjC domain containing demethylase. Notably, bacteria do not have KDM1 homologs^54^. These data indicate that geldanamycin has an evolutionary selectivity to JmjC KDMs. While the structural basis for selectivity of geldanamycin and its analogs against KDM remains to be solved, the selectivity is probably determined by the differences of 17-position of benzoquinone ring (17-methoxy group of geldanamycin, 17-dimethylaminoethylamino of 17-DMAG, and 17-N-allylamino of 17-AAG). Based on structural studies, this position is highly solvent exposed in the Hsp90-Geldanamycin crystal complex and is a poor candidate for additional Hsp90 contacts^45^, suggesting that it is not essential to Hsp90 binding.

In summary, we identified Hsp90 inhibitors, geldanamycin and its analog 17-DMAG, as novel and potent KDM inhibitors. We also found that PAX3-FOXO1 is a Hsp90 client, which was destabilized by geldanamycin. Our findings support a concept that PAX3-FOXO1 creates an epigenetic dependency to KDMs and chaperone dependency to Hsp90, and thus dually targeting KDMs and Hsp90 is a potentially valuable therapeutic option for PAX3-FOXO1-driven aRMS (**Supplementary Fig. 7**). However, more side-effects might be also expected. Nevertheless, in addition to inhibiting Hsp90 and KDMs, the therapeutic efficacy of 17-DMAG could also be due to other off-targeting molecules.

## Materials and Methods

### Cell lines and reagents

#### Cell lines

293T, U2OS, Rh30, Rh41 cell lines were purchased from ATCC and validated by short tandem repeat (STR) using Promega PowerPlex 16 HS System once per month. PCR-based method was used for detection of Mycoplasma with LookOut Mycoplasma PCR Detection Kit (Sigma) and JumpStart *Taq* DNA Polymerase (Sigma) once per month to ensure cells were mycoplasma negative.

#### Compounds

17-DMAG, 17-AAG and geldanamycin were purchased from Selleckchem. MG132 and Ciclopirox were purchased from Sigma. Vincristine, Irinotecan, and JQ1 were purchased from MedChem Express (MCE).

#### Antibodies

The anti-PAX3-FOXO1 mouse monoclonal antibody was provided by Dr. Liang Cao (NCI). The anti-KDM4B antibody (A301-478A) was purchased from Bethyl Laboratories. The anti-Actin antibody (A2228) was purchased from Sigma. Anti-H3K4me3 (07-473) and Anti-normal mouse IgG antibody (12-371) was purchased from Millipore. The anti-total H3 (4499), anti-H3K9me3 (13969), anti-H3K36me3 (9763), anti-H3K4me3 (9751) antibodies were purchased from Cell Signaling Technology (CST). Anti-FOXO1 antibody (2880) that recognizes the fusion protein PAX3-FOXO1 was purchased from CST. Anti-Hsp90 antibody (F-8) (sc-13119), anti-Hsp70 antibody (C92F3A-5)(sc-66048) and p53 antibody (DO-1) were purchased from Santa Cruz Biotechnology. Secondary horseradish peroxidase(HRP)-conjugated goat anti-mouse (31430) and goat anti-rabbit (31460) antibodies were purchased from Thermo Fisher Scientific.

#### Plasmids

MSCV-KDM4B(wt)-RFP and MSCV-KDM4B(H189A)-RFP constructs were generated by PCR of the full length of wild type and mutant KDM4B following by subcloning into MSCV-IRES-RFP plasmid, and standard retroviral packaging. U2OS cells were transduced with retroviral particles for high-content image screen. pCMV-HA-KDM4B was obtained from Addgene (24181). The catalytic domain of KDM4B(1-348) was subcloned into pET28a to produce histidine tagged KDM4B protein by Protein Production Facility at St Jude for TR-FRET and MALDI-FTICR screening.

### TR-FRET demethylation functional assay

Stock compound solutions (10 mM compound in DMSO) or DMSO only (vehicle control) were transferred to the individual wells in low volume black 384-well assay plates containing 1.5 µM biotin-H3K9me3 in 10 µL assay buffer [50 mM Tris-HCl (pH 8.0), 1 mM α-ketoglutarate, 80 μM FeSO_4_, 2 mM ascorbic acid, 0.01% BSA] by using a V&P 384-well pintool (V&P Scientific, San Diego, CA) at 30 nL/well. KDM4B protein (750 nM) or buffer only was then dispensed (5 µL/well). After a brief spin down and shake, the plates were incubated at room temperature for 30 min. Detection reagent (5 µL/well) of 8 nM Tb-anti-H3K9me2 antibody and 8 nM AF488-streptavidin was dispensed, followed by a brief spin down, shake and 15 min room temperature incubation. The TR-FRET signal (fluorescence emission ratio of 10,000 × 520 nm/490 nm) from each well was collected with a PHERAstar *FS* plate reader (BMG LABTECH Inc., Cary, NC). The final tested compound concentration was 20 µM and the final DMSO concentration was 0.2% for all wells in the primary screening. The DMSO control wells with KDM4B protein and those without KDM4B protein were used as negative (0% inhibition) and positive (100% inhibition) controls, respectively. The individual compound activities were normalized to those of negative and positive controls. Compounds with %Inhibition ≥ 30% were selected as hits for DR analysis (10 concentrations, following a 1:3 serial dilution scheme; final concentration range 4.7 nM to 93.3 µM, in triplicates) under similar assay condition as the primary screening, with the exception of the final DMSO concentration at 0.93% for all assay wells. The activity data for individual chemicals were normalized to that of positive and negative controls and fit into sigmoidal DR equation, if applicable, to derive DR curves and IC_50_ values with GraphPad Prism 8.0.

### High-content immunofluorescence imaging assay

1000 U2OS-KDM4B expressing cells in 25 μl of media were plated into each well of a poly-D-lysine coated Perkin Elmer 384-well View plates (Perkin Elmer 6007710) with a Thermo Scientific Wellmate. The cells were then grown for 18 hours overnight before they were drugged using a VP scientific pintool with S100 pins. The cells were then treated with compound for a twenty-four hours. Following treatment, the cells were fixed with 4% formaldehyde for 20 minutes at 37°C and permeabilized with 0.1% Triton-X 100 for 15 minutes at 25°C. Fixative was removed and each well washed with PBS. Cells were blocked using 1% BSA in PBS for 1 hour at 25°C. The primary antibody against trimethyl-histone H3 at Lys9 (Millipore 07-442) was used at 1/400 dilution in 1% BSA in PBS. This mixture was added to each well before incubation overnight at 4°C. Each well was then washed 3 times with PBS using a Biotek plate washer, and incubated for 1 hour at 25°C with a solution containing 1/400 goat α-rabbit-Alexa-488 (Cell Signaling 4412S) and 1 μM Hoechst 34580 to detect nuclear material (H21486 Molecular Probes.) Two images were captured of each well at 10X using a GE Healthcare InCell 6000 at 405 to detect nuclear staining and 488 nM to detect H3K9me3. The number of nuclear objects in each well, as detected through Hoechst staining, was compared to the number of cells in each well expressing a minimum amount of H3K9me3 as determined by Alexa-488 fluorescence (1.5 million counts total intensity), to identify the percentage of cells in each considered “H3K9me3 Positive.” Averages shown are the result of eight replicate measurements per data point.

### MALDI-FTICR mass spectrometry-based demethylation assay

To assess the inhibition potency of geldanamycin and its analogs, we used a MALDI-TOF MS-based platform developed by our group. A truncated version of KDM4B that contains only the JmjC catalytic domain, KDM4B(1-348) was used. KDM4B(1-348), to a final concentration of 250 nM (in 50 mM Tris base, pH 7.3), was incubated with different concentrations of each compound (10 concentrations in total) for 1 hour at room temperature. The final concentration of DMSO in each well was 1 %. As negative control, KDM4B(1-348) was incubated with 1 % DMSO. Positive control wells contained 10 % formic acid. Reactions were initiated upon adding the “substrate mixture” (200 μM α-ketoglutarate, 100 μM ascorbate, 10 μM NH4Fe(SO4)2, and 10 μM H3K9me3(1-15) peptide) to each well. Reactions were incubated for 90 min at room temperature (to achieve ∼ 20 % Turnover) prior to quenching with 10 % formic acid. Assays were performed in triplicates (n = 3).

Two microliters from each well were mixed with 18 μL of MALDI matrix solution (20 mg/mL of 2,5 dihydroxybenzoic acid dissolved in 95 % methanol), from which 1 μL was spotted on a 384 AnchorChip® MALDI target plate. Crystalized samples were then analyzed using a 7 T Solarix XR Fourier Transform Ion Cyclotron Resonance (FT-ICR) mass spectrometer (Bruker Co., MA, USA). The MALDI-FTICR parameters were optimized as follows: laser power = 20 % at 200 shots, laser shot frequency = 800 Hz, isolated Q1 m/z = 1530 ± 20. The mass of two species were detected: m/z = 1530.87 (substrate), and m/z = 1516.85 (dimethylated product). The following formula was used to calculate % Turnover: 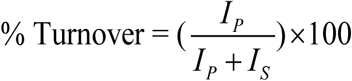

Where IP and IS are the ion intensities of product and substrate, respectively. Values for % Turnover were then normalized based on negative and positive controls.

### ALPHA Screen demethylation assay

All reagents were provided by BPS company. All of the enzymatic reactions were conducted in duplicate at room temperature for 60 minutes in a 10 µl mixture containing assay buffer, histone H3 peptide substrate, demethylase enzyme, and the test compounds.

These 10 µl reactions were carried out in wells of 384-well Optiplate (PerkinElmer). The dilution of the compounds was first performed in 100% DMSO with the highest concentration at 0.5mM. Each intermediate compound dilution (in 100% DMSO) will then get directly diluted 30x fold into assay buffer for 3.3x conc (DMSO). Enzyme only and blank only wells have a final DMSO concentration of 1%. From this intermediate step, 3 µl of compound is added to 4 µl of demethylase enzyme dilution is incubated for 30 minutes at room temperature. After this incubation, 3 µl of peptide substrate is added. The final DMSO concentration is 1%. After enzymatic reactions, 5 µl of anti-Mouse Acceptor beads (PerkinElmer, diluted 1:500 with 1x detection buffer) or 5 µl of anti-Rabbit Acceptor beads (PerkinElmer, diluted 1:500 with 1x detection buffer) and 5 µl of Primary antibody (BPS, diluted 1:200 with 1x detection buffer) were added to the reaction mix. After brief shaking, plate was incubated for 30 minutes. Finally, 10 µl of AlphaScreen Streptavidin-conjugated donor beads (Perkin, diluted 1:125 with 1x detection buffer) were added. In 30 minutes, the samples were measured in AlphaScreen microplate reader (EnSpire Alpha 2390 Multilabel Reader, PerkinElmer). Enzyme activity assays were performed in duplicates at each concentration. The A-screen intensity data were analyzed and compared. In the absence of the compound, the intensity (Ce) in each data set was defined as 100% activity. In the absence of enzyme, the intensity (C0) in each data set was defined as 0% activity. The percent activity in the presence of each compound was calculated according to the following equation: %activity = (C-C0)/(Ce-C0), where C= the A-screen intensity in the presence of the compound.

The values of % activity versus a series of compound concentrations were plotted using non-linear regression analysis of Sigmoidal dose-response curve generated with the equation Y=B+(T-B)/1+10((LogEC50-X)×Hill Slope), where Y=percent activity, B=minimum percent activity, T=maximum percent activity, X= logarithm of compound and Hill Slope=slope factor or Hill coefficient. The IC50 value was determined by the concentration causing a half-maximal percent activity.

### Microscale Thermophoresis assay

KDM4B was labeled with a cysteine reactive fluorescent dye (Protein Labeling Kit RED-MALEIMIDE, Nanotemper GmbH). A serial dilution of titrant was prepared in MST buffer containing 10 mM HEPES (pH 7.5), 200 mM NaCl, 2 mM NiCl2, 5% DMSO and 0.05% Tween 20. An equal volume of diluted titrant and the constant concentration of labeled KDM4B were added and loaded in a standard treated capillaries (Nanotemper GmbH). Binding measurements were performed on a Monolith NT.115 Blue/Red instrument (Nanotemper GmbH) at 40 % LED power and 40 % MST power. The data were analyzed using MO Affinity Analysis software (Nanotemper GmbH).

### Phylogenetic tree analysis

Amino acid sequences of KDM domains of histone lysine demethylases were aligned with Clustal Omega program (https://www.ebi.ac.uk/Tools/msa/clustalo/) and the phylogenetic tree was generated using neighbor-joining method, which was shown using iTOL program (https://itol.embl.de).

### RNA-seq

Total RNA was extracted from xenograft tissues by RNeasy Mini Kit (cat. # 74104) from QIAGEN. Paired-end sequencing was performed using the High-Seq platform with 100bp read length. Reads were aligned to the human GRCh37-lite using SJCRH’s Strongarm pipeline. Counts per gene were obtained using htseq-count version 0.6.1 with Gencode vM5 level 1and 2 gene annotations. Counts were normalized with VOOM and analyzed with LIMMA within the R statistical environment. Significance was defined as having a false discovery rate (FDR) <0.05. VOOM normalized counts were analyzed with Gene Set Enrichment Analysis (GSEA)^55^.

### Western blot

Cells were washed twice with ice-cold phosphate-buffered saline (PBS) and then directly lysed on ice with 2X sample loading buffer (0.1 M Tris HCl [pH 6.8], 200 mM dithiothreitol [DTT], 0.01% bromphenol blue, 4% sodium dodecyl sulfate [SDS], 20% glycerol). On ice, cell lysates were briefly sonicated once for 5 seconds at 40% amplitude output followed by 25 minutes heating at 95 °C. Afterwards, cell lysates were briefly centrifuged at 13,000 × g at room temperature for 1 minute. Then, 20 µl of cell lysates were separated on 4-12% tris-glycine SDS-polyacrylamide gel electrophoresis (SDS-PAGE) from Invitrogen, and transferred to methanol-soaked polyvinylidene difluoride (PVDF) membranes from Millipore. Membranes were blocked in PBS buffer supplemented with 0.1% TWEEN 20 and 5% skim milk (PBS-T), and incubated for 1 hour at room temperature under gentle horizontal shaking. Afterwards, membranes were incubated overnight at 4°C with the primary antibodies under gentle horizontal shaking. The primary antibodies were prepared in PBS-T with the following dilutions: anti-KDM4B (1:1000), anti-PAX3-FOXO1 (1:200), anti-FOXO1 (1:1000), anti-Hsp90 (1:1,000), anti-actin (1:5,000), anti-total H3 (1:2,000), anti-H3K4me3 (1:2,000), anti-H3K9me3 (1:2,000) and anti-H3K36me3 (1:2,000). Next day, membranes were washed 3 times (each wash for 5 minutes) with PBS-T at room temperature. Protected from light, membranes were then incubated with goat anti-mouse or goat anti-rabbit HRP-conjugated secondary antibodies (1:5,000) for 1 hour at room temperature. Then, membranes were washed 3 times (each wash for 5 minutes) with PBS-T at room temperature. Lastly, membranes were incubated for 1 minute at room temperature with SuperSignal West Pico PLUS Chemiluminescent Substrate (34580, Thermo Fisher Scientific), and the bound antigen-antibody complexes were visualized using Odyssey Fc Imaging System (LI-COR Corp., Lincoln, NE).

### Immunoprecipitation

Cells were washed twice with ice-cold PBS and then directly lysed on ice with co immunoprecipitation (co-IP) buffer (25 mM Tris-HCl [pH 7.5], 150 mM NaCl, 1 mM ethylenediaminetetraacetic acid [EDTA], 1% nonyl phenoxypolyethoxylethanol [NP40], 5% glycerol) supplemented with phosphatase (Roche) and protease (Roche) inhibitor cocktails. The cell lysate was transferred to a 2-ml Eppendorf tube, and incubated on ice for 15 minutes, and vortexed every 5 minutes. Then, the cell lysate was centrifuged at 13,000 × g at 4 °C for 15 minutes. The pre-cleared supernatant was incubated with rotation at 4 °C overnight with 4 µg of anti-HSP90 and 4 µg of anti-normal IgG as a negative control. Next day, 50 µl of protein A/G magnetic beads (88802, Thermo Scientific Fisher) were washed 3 times at room temperature with the co-IP buffer, and then added to each pre-cleared supernatant for 1 hour incubation with rotation at 4 °C. Afterwards, the supernatant (flow-through) was discarded; the beads were washed 3 times with co-IP buffer, eluted with 50 µl of the 2X sample loading buffer and heated for 10 mins at 95 °C. Input lysate was heated for 25 mins at 95 °C. 20 µl of co-IP and 20 µl of input reactions were subjected to SDS-PAGE and immunoblotting with anti-Hsp90 and anti-PAX3-FOXO1 antibodies (as described above).

### Immunohistochemistry

Xenografts were fixed in 10% neutral buffered formalin, embedded in paraffin, sectioned at 4 μm, stained with hematoxylin and eosin and reviewed by light microscopy using an upright Nikon Eclipse Ni microscope (Nikon Instruments, Inc.). Immunohistochemistry was performed on 4 μm thick tissue sections mounted on positively charged glass slides (Superfrost Plus; Thermo Fisher Scientific, Waltham, MA), and dried at 60°C for 20 minutes. The procedures for immunohistochemistry were performed using a Ventana DISCOVERY ULTRA autostainer (Roche). Heat induced epitope retrieval was applied for 1 hour using cell conditioning 1 buffer (CC1, Roche, #950-500) followed by the application of anti-CD31 (Histobiotec, DIA-310, 1:50) or anti-Cleaved Caspase 3 (BioCare Medical, CP229C, 1:500) for 32 minutes. The following reagents were used for visualization: DISCOVERY OmniMap anti-Rat HRP (Roche,760-4457) for CD31 or OmniMap anti-Rabbit HRP (Roche, 760-4311) and the DISCOVERY ChromoMap DAB kit (Roche, 760-159), which was applied for 8 min at room temperature. Tissues were counterstained with Hematoxylin II (Roche, 790-228) for 12 min and Bluing reagent (Roche, 760-2037) for 4 min as a post-counterstain procedure. TUNEL was performed using the *In situ* Cell Death Detection Kit (Roche, 11684817910) according to the manufacturer’s instructions. The quantification of Caspase 3, TUNEL and CD31 was performed using ImageJ IHC tool box (for Caspase 3 and TUNEL) and Vessel Analysis plug in program (for CD31), and unpaired student t test was used to compare the difference between vehicle and treatment.

### Animal experiments

All murine experiments were done in accordance with a protocol approved by the Institutional Animal Care and Use Committee of St. Jude Children’s Research Hospital. Subcutaneous xenografts in Figure 5 and Figure 6E were established in CB17 severe combined immunodeficient mice (CB17 *scid*, Taconic) by implanting 5×10^6^ cells in Matrigel. Subcutaneous xenografts in Figure 6A - 6D were established NOD.Cg-*Prkdc*^*scid*^*Il2rg*^*tm1Wjl*^/SzJ (NOD *scid* gamma, NSG) mice by implanting 5×10^6^ cells in Matrigel. Tumor measurements were done weekly using electronic calipers, and volumes calculated as width π/6 × *d*^3^ where *d* is the mean of two diameters taken at right angles. Subcutaneous xenografts were treated with 25 mg/kg of 17-DMAG or 50 mg/kg of 17-AAG via intraperitoneal injection twice daily, every four days. 17-DMAG was dissolved in 1% DMSO, 1% TWEEN^®^ 80 (#P4780 from Sigma), 30% PEG300 (#202371 from Sigma) and 68% ddH_2_O. 17-AAG was dissolved in 5% DMSO and 95% corn oil. Vincristine was administered in a dose of 0.38 mg/kg via IP injection once daily every week. Vincristine was dissolved in 100% saline. Irinotecan was administered in a dose of 1.25 mg/kg via IP injection once daily, for 5 days on and 2 days off schedule. Irinotecan was dissolved in 5% DMSO and 95% saline. Mice were sacrificed because of an adverse event before they had completed 14d and were removed from the data set. Tumor response: For individual mice, progressive disease (PD) was defined as < 50% regression from initial volume during the study period and > 25% increase in initial volume at the end of study period. Stable disease (SD) was defined as < 50% regression from initial volume during the study period and ≤ 25% increase in initial volume at the end of the study. Partial response (PR) was defined as a tumor volume regression ≥50% for at least one time point but with measurable tumor (≥ 0.10 cm^3^). Complete response (CR) was defined as a disappearance of measurable tumor mass (< 0.10 cm^3^) for at least one time point.

### Statistical analysis

To determine statistical significance, the unpaired, two-tailed Student *t* test was calculated using the *t* test calculator available on GraphPad Prism 8.0 software. A *p* value of less than 0.05 was considered statistically significant. Kaplan-Meier survival analysis was calculated using log-rank (Mantel-Cox) method in GraphPad Prism 8.0 software.

The Kruskal Wallis test was utilized to determine if there was a statistically significant different among the 4 treatment groups at each time point. The exact Wilcoxon Rank Sum test was utilized to determine if there was a statistically significant difference between receiving one treatment vs receiving the combination treatment. All statistical analyses were conducted in SAS 9.4 and a two-sided significance level of *p*<0.05 was determined a priori.

## Acknowledgments

This work was partly supported by American Cancer Society-Research Scholar (130421-RSG-17-071-01-TBG, J.Y.) and National Cancer Institute (R03CA212802-01A1, J.Y.; 1R01CA229739-01, J.Y., T.C., S.W., A.D.; R35-GM118041, T.C.),

## Contributions

A.A.Z, S.S., W.L., J.L., J.F., A.A., B.C., J.B., M.K.Y., A.A.T, D.F., J.M., H.T., B.Y., S.D., and Z.L. performed research. A.A.Z, D.C., S.S., W.L., J.L., A.A., R.L., M.K.Y., J.Y. P.B., Z.L. analyzed data. J.Y., A.M.D. T.C., S.W., and Z.R. designed the research. J.Y. wrote the paper with input from C.T., A.M.D.

